# Examining infantile facial features and their influence on caretaking behaviors in free-ranging Japanese macaques (*Macaca fuscata*)

**DOI:** 10.1101/2023.10.12.562000

**Authors:** Toshiki Minami, Takeshi Furuichi

**Author notes:** Corresponding author: ORCID: 0000-0001-5476-3896.

## Abstract

Facial features of immature individuals play a pivotal role in eliciting caretaking behaviors in humans. It has been posited that non-human animals share particular infantile facial features with humans, which can elicit caregivers’ attention and caretaking behaviors. Nevertheless, the empirical examination of this hypothesis is extremely limited. In this study, we investigated infantile facial features in Japanese macaques (*Macaca fuscata*), their developmental processes, and their correlation with caretaking and infant behaviors, based on 470 facial photographs from one free-ranging group. We measured the size of facial parts and evaluated infantile facial features in Japanese macaques with non-contact procedures with the animals. The results indicated that, although some partial species differences were observed, the infantile facial features in Japanese macaques were broadly consistent with those previously observed in humans and great apes. Furthermore, half of the infant subjects displayed non-linear developmental trajectories of infantile faces similar to those suggested in humans. These changes were sex-based, with most females demonstrating a linear trajectory and most males a hump-shaped development in their facial proportions. However, unlike previous studies in humans, infantile faces were not significantly associated with maternal or non-maternal caretaking behaviors, nor were their developmental changes correlated with infant behavioral development. These findings indicate that while many aspects of infantile facial features are shared among particular primates, humans may have evolved a uniquely elevated preference for selecting such features among the primate lineage.

## Introduction

Primates, including humans, exhibit extended periods of immaturity [1], and neonates face survival challenges when deprived of continuous and adequate support. Moreover, the care received during immaturity has long-term effects on development [2]. For example, mortality rates of Japanese macaque (*Macaca fuscata*) infants without sufficient maternal care are higher [3]. Additionally, early postnatal affiliation with caregivers significantly affects infant development in primates [4, 5]. Thus, studies on primate caretaking behaviors are crucial for understanding the factors affecting the survival and maturation of immature individuals.

In humans, one of the key triggers for caretaking behaviors is the physical features of immature individuals [6]. Lorenz [7] proposed that infants possess distinct physical features that stimulate their care. While his suggestion encompassed overall morphological characteristics, subsequent studies have highlighted specific facial features, such as a greater facial width relative to head length (i.e., a rounded head), a longer forehead relative to the facial length (i.e., low-set eyes), larger eyes relative to the facial width, and a smaller nose and mouth relative to the facial size [8]. Experimentally enhancing these features when evaluating infant facial stimuli can increase infant attractiveness and capture and retain the observer’s attention [8−16]. Furthermore, watching infant faces can activate neural reward and motivational systems [17] and motivate caretaking behavior in recipients [8]. Although studies on the link between facial features and caretaking behaviors are restricted, Langlois et al. [18] reported that neonates rated as more attractive by non-caregivers experienced more intimate interactions with their caregiving mothers. These findings suggest that the facial appearance of human infants can promote positive responses from caregivers and potentially elicit tangible caretaking behaviors.

Contrary to our intuition, the prominence of these facial features may not always be more pronounced in less mature infants. For example, when compared to preterm infants of the same age, full-term infants have more emphasized facial features such as forehead length, facial width, and eye width [19]. Furthermore, the development of facial cuteness in infants follows a hump-shaped trajectory with age, peaking between 6 and 11 months of age, rather than immediately postpartum [15, 20, 21, 22]. This trend is intriguing, given that newborns are more vulnerable [20]. As a potential explanation for this non-linear developmental change, Negayama [15] proposed a link to infant mobility. The peak age of human infantile facial attractiveness coincides with the period when they begin exploring their environment independent of their caregivers, thereby increasing the likelihood of encountering potential dangers. The most attractive faces at this developmental stage could more strongly capture the attention of caregivers and help mitigate heightened danger. Although it is necessary to examine this hypothesis empirically, the development of infantile features may also have an evolutionary origin that captures elevated attention from caregivers.

Lorenz [7] proposed that infantile morphologies and the increased attention paid to them are widespread across various animal species, a long-acknowledged perspective. However, empirical investigations on non-human animals have only recently begun. For instance, Kawaguchi et al. [23] remains the only study to examine the interspecific similarities and differences in infantile facial features among primates. They compared the facial morphologies of great apes, including humans, and revealed the following conserved facial features in infants: expanded facial width (round and short faces), a prolonged forehead (eyes positioned lower on the face), enlarged eyes, and an inverted triangular configuration. In contrast, the smaller nose and mouth, which are typical characteristics of human infants, were not shared among the great apes as infantile features [23]. To advance our understanding of cross-species similarities and variations in infantile facial features, an examination across species beyond the great apes is necessary.

Research is also underway to explore how non-human primates react to the faces of conspecific infants. By presenting infant and adult images, preference for infant faces has been documented in chimpanzees (*Pan troglodytes*), Barbary macaques (*Macaca sylvanus*), and rhesus macaques (*Macaca mulatta*) [24−26]. However, their reactions were likely to be motivated by infantile facial color rather than facial shape. Additionally, bonobos (*Pan paniscus*) and Japanese macaques did not exhibit a significant preference for infant faces [27, 28]. These investigations suggest that non-human primates are less attracted to infantile faces, while humans strongly prefer infantile facial morphology. However, it is unclear whether infantile facial features are linked to the recipient’s motivation for caretaking, which can directly contribute to infant survival and development. To test Lorenz’s [7] hypothesis in primates, a novel study on the direct association between infant facial appearance and caretaking behavior is essential.

Whether the hump-shaped development of infantile faces observed in humans is also present in non-human primates remains uncertain. In the only relevant study, Sanefuji et al. [22] presented images of infant chimpanzees, rabbits (*Oryctolagus cuniculus*), dogs (*Canis familiaris*), and cats (*Felis syvestris catus*) to college students. The findings revealed that the most attractive faces across all species were not those of neonates, but rather of infants several months old. Humans perceive the attractiveness of heterospecific faces using the same criteria as they do for conspecific infants [29, 30]. Hence, if infantile facial features are shared across animal taxa [7], this suggests the potential existence of non-linear development of infantile faces in non-human species. Additionally, similar to humans, non-human primate infants would face escalated risks of injury when they begin independent exploratory behaviors. Therefore, if infantile faces indeed facilitate caretaking behaviors, non-human infants might benefit from having pronounced facial features during this critical period of their infancy [15].

To further our understanding of the associations between infantile facial features and caretaking behaviors in non-human primates, this study focused on Japanese macaques, a species unrelated to great apes, to investigate the following four questions: (1) Do Japanese macaques exhibit facial features similar to those of great apes, including humans? [7] (2) Is there a correlation between the physical proportions of infantile faces and caretaking behaviors among infant Japanese macaques? [7] (3) Do the facial proportions of infant Japanese macaques show a hump-shaped development, akin to human infants? [15, 20, 21, 22] (4) Is there a clear association between pronounced infantile faces and the initiation of exploratory behavior in Japanese macaque infants? [15] To answer these questions, we collected 470 facial photographs from nearly all individuals within a single group of Japanese macaques, quantitatively evaluated their facial features, and examined the development of infantile faces and their potential associations with behavioral patterns.

## Methods

### Study site and subjects

This study was conducted at Arashiyama Monkey Park Iwatayama, Kyoto, Japan. This private park has consistently fed one group of free-ranging Japanese macaques (Arashiyama group) since 1954, presenting invaluable opportunities for visitors to observe wild monkeys [31]. Throughout this study, the Arashiyama group consistently comprised approximately 130 monkeys. These macaques inhabit their native habitats, with unrestricted access to natural food resources and freedom in their behavior. The park has maintained genealogical records of these monkeys since the 1950s [31], availing accurate information on individual birthdays and ages.

The present study included 128 Japanese macaques, comprising 65 adult females and five adult males (≥ 5 years old), 25 juvenile females and 25 juvenile males (1−4 years old), and three infant females and three infant males (<1 year old). All targets were monkeys of known age affiliated with this group in 2021. Table 1 provides detailed information on the infant subjects.

**Table 1.**
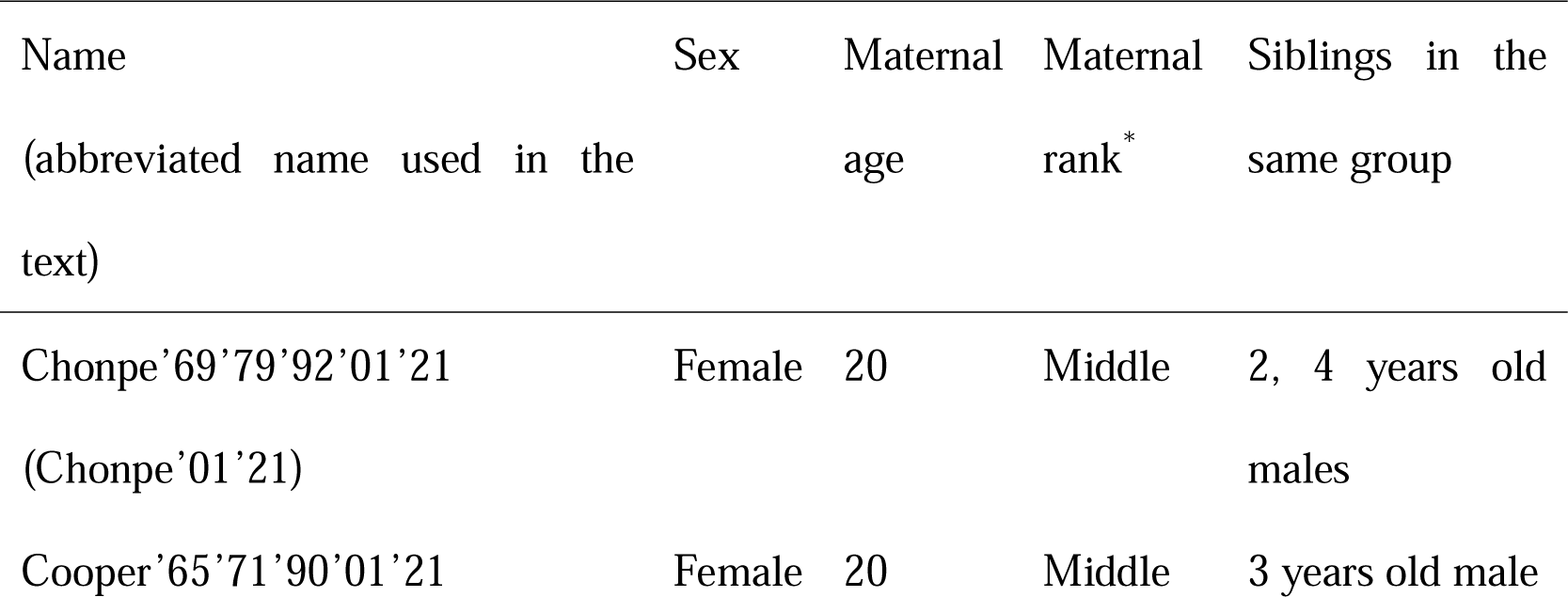
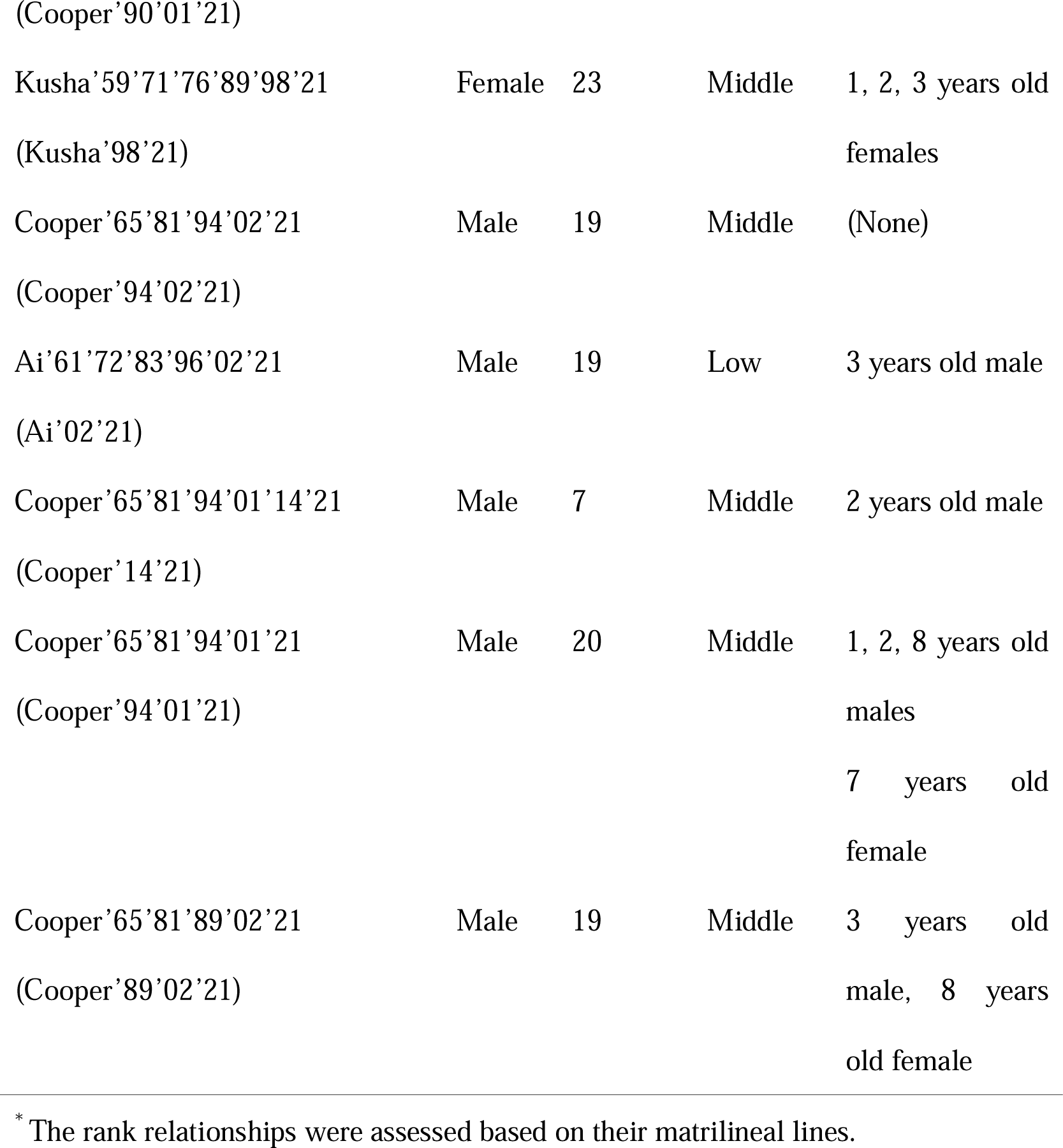
Individual information on the eight Japanese macaque infants whose facial development and behavior were monitored.

We chose the Arashiyama group for this study for four reasons. First, to study primate species beyond the great apes, the Japanese macaque was one of the apt candidates because this species falls in the tribe Catarrhini together with the apes [32]. Second, the extensive efforts of the park to maintain detailed genealogical records provided us with comprehensive information on the birthdates and ages of all the subjects. This information is crucial for precise developmental studies. Third, this study required numerous facial photographs to examine temporal changes in facial development in detail. The Arashiyama group stayed in the feeding area with minimal obstruction during much of the day and was habituated to human observers, enabling us to collect an adequate number of images. Fourth, we selected subject animals within their native habitats rather than in captivity to ascertain the associations between infant faces and unrestricted behavior. As such, the Arashiyama group was an ideal subject for this study.

### Morphometrics

We used a non-contact technique to measure the monkey faces, which assesses the proportional size of facial features based on frontal facial photographs [8, 33, 34]. This methodology facilitated data acquisition while minimizing potential stress on the subjects. The analysis focused on six facial features: face width relative to head length (FWHL), forehead length relative to face length (FoLFaL), eye width relative to face width (EWFW), nose length relative to head length (NLHL), nose width relative to face width (NWFW), and mouth width relative to face width (MWFW). These facial features have been regarded as typical infantile features in humans [8] and were discussed in a cross-species study [23]. Therefore, our methodology allowed for comparisons between the results from Japanese macaques and previous knowledge from other primates.

We collected facial images for 141 days, from December 2020 to August 2022. Notably, there was a brief intermission in data collection from April 25 to May 21, 2021, owing to the temporary closure of the park caused by the COVID-19 outbreak. To capture the photographic data, we used a digital camera (Canon EOS M6 with Canon EF–S55–250 mm F4–5.6 IS Stem lens) or a video camera (Panasonic HC-WZ590M-W) positioned at a distance exceeding 2 m from the subjects. Considering the focus on infant development, the frequency of capturing facial photographs differed according to the participant’s age. Adults were photographed once throughout the study period. Juveniles underwent photography approximately every three months from April 2021 to August 2022, with a maximum of four images per subject. For infants, images were captured weekly from birth to 24 weeks, followed by a transition to monthly intervals until the age of 1 year, yielding a maximum of 31 photographs per infant. In total, we captured 470 facial photographs: 70 from adults, 184 from juveniles, and 216 from infants.

GIMP (version 2.10.22: https://www.gimp.org/) was used for photographic analysis. All photos were cropped into a square format with dimensions of 500 × 500 pixels, which included the entire faces of the subjects. Subsequently, the first author manually plotted 15 landmarks on each photo (Fig 1 and Table 2). Following this process, the lengths of nine distinct facial parts (outlined in Table 3) were quantified in terms of pixels using the measure tool in GIMP. Finally, six target indices (FWHL, FoLFaL, EWFW, NLHL, NWFW, and MWFW; Table 4) were calculated.

**Fig 1.**
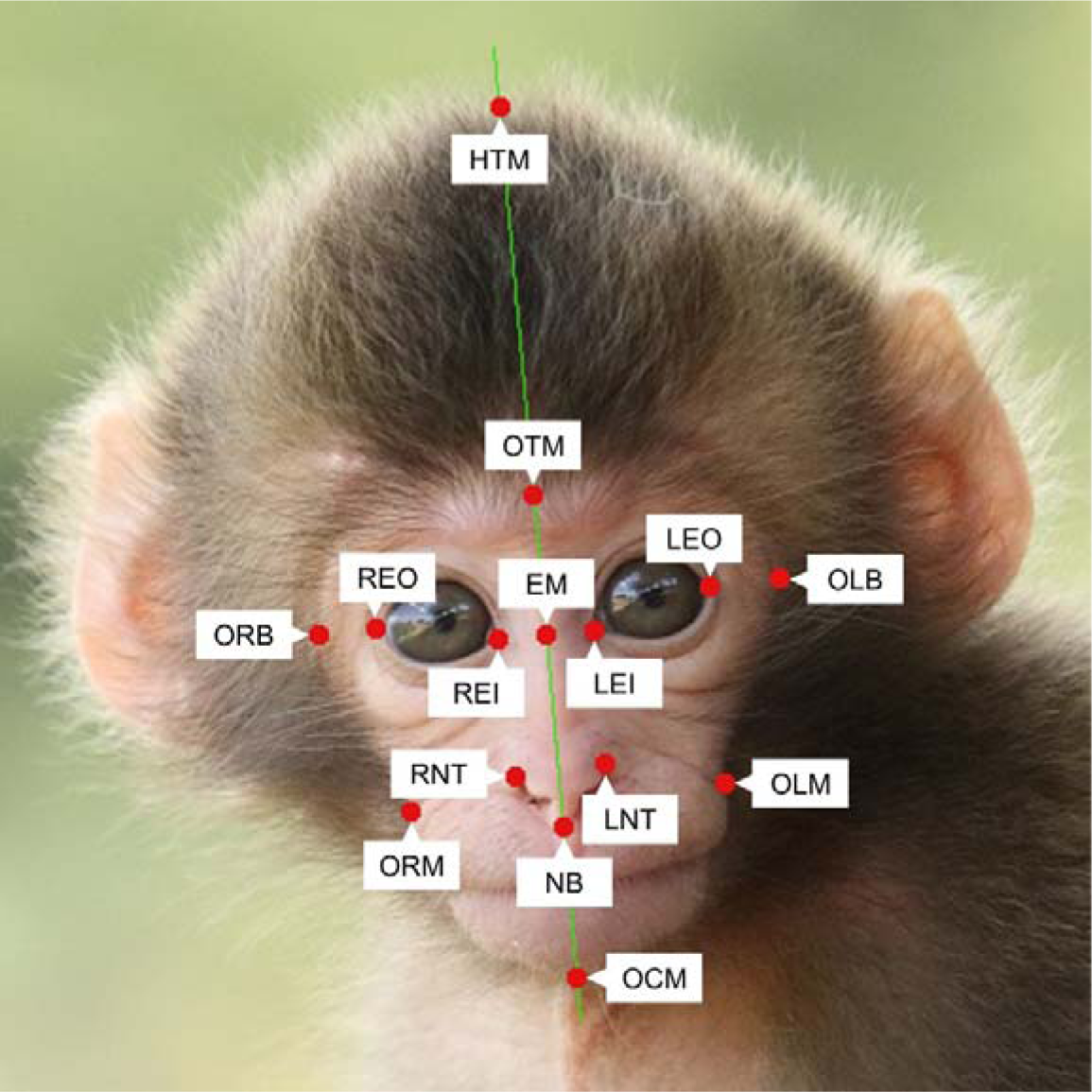
Example of landmark plots on the frontal face view of Japanese macaque infants.

**Table 2.** Definitions of Japanese macaque infant facial landmarks evaluated in this study.

| Landmark | Abbreviation | Definition |
| --- | --- | --- |
| HeadTop_Mid | HTM | Centre of top of the head |
| OutlineTop_Mid | OTM | Top center of the outline of the face |
| OutlineRight_Brow | ORB | Outline at right point horizontally aligned to eye outers |
| OutlineLeft_Brow | OLB | Outline at left point horizontally aligned to eye outers |
| OutlineChin_Mid | OCM | Point on the outline of the center of the chin |
| Eyes_Midpoint | EM | Point halfway between two eye inners on the midline of the face |
| RightEye_Inner | REI | The inner corner of the right eye |
| RightEye_Outer | REO | The outer corner of the right eye |
| LeftEye_Inner | LEI | The inner corner of the left eye |
| LeftEye_Outer | LEO | The outer corner of the left eye |
| OutlineRight_Mouth | ORM | The right corner of the mouth |
| OutlineLeft_Mouth | OLM | The left corner of the mouth |
| RightNostril_Top | RNT | Center of the top edge of the right nostril |
| LeftNostril_Top | LNT | Center of the top edge of the left nostril |
| Nostrils_Bottom | NB | Point halfway between the bottoms of two nostrils |

The definitions were taken from the “primate\_face” model of DeepLabCut Model Zoo (http://www.mackenziemathislab.org/dlc-modelzoo), and only some of the expressions were changed.
“Outline” means the edge of the hairless area.

**Table 3.** Facial parts and inter-rater reliability of the measurements.

| Facial part | Abbreviation | Definition<sup>1</sup> | ICC (3,1) | 95% CI | Assessment<sup>2</sup> |
| --- | --- | --- | --- | --- | --- |
| Head length | HL | HTM-OCM | 0.89 | 0.85-0.93 | Good-Excellent |
| Forehead length | FoL | HTM-OTM | 0.98 | 0.98-0.99 | Excellent |
| Face length | FaL | OTM-OCM | 0.95 | 0.93-0.97 | Excellent |
| Face width | FW | ORM-OLM | 0.81 | 0.74–0.87 | Moderate–Good |
| Right-eye width | REW | REI-REO | 0.91 | 0.88–0.94 | Good–Excellent |
| Left-eye width | LEW | LEI-LEO | 0.90 | 0.86–0.93 | Good–Excellent |
| Nose length | NL | EM-NB | 0.97 | 0.96–0.98 | Excellent |
| Nose width | NW | RNT-LNT | 0.73 | 0.64–0.81 | Moderate–Good |
| Mouth width | MW | ORM-OLM | 0.83 | 0.76–0.88 | Good |

<sup>1</sup> Distance between two facial landmarks
<sup>2</sup> The reliability was assessed based on the guideline proposed by Koo and Li (2016): “Excellent” for 95% CI > 0.90, “Good” for 0.75 < 95% CI < 0.90, and “Moderate” for 0.50 < 95% CI < 0.75.

**Table 4.** Summary of the six facial measurement indices calculated for analysis of the facial development in Japanese macaque infants.

| Index | Abbreviation | Formula |
| --- | --- | --- |
| Relative size of face width | FWHL | FW/HL |
| Relative size of forehead length | FoLFaL | FoL/FaL |
| Relative size of eye width | EWFW | ((REW + LEW)/2)/FW |
| Relative size of nose length | NLHL | NL/HL |
| Relative size of nose width | NWFW | NW/FW |
| Relative size of mouth width | MWFW | MW/FW |

To ascertain the reliability of the measurements performed by the first author, an additional rater, who was blinded to the study aims and had limited familiarity with Japanese macaques, placed landmarks on a subset of 118 images (25.1% of the total), including 18 adults, 46 juveniles, and 54 infants. Using these landmarks, we computed nine facial parts (Table 3) and evaluated their consistency with the measurements obtained by the first author using the intraclass correlation coefficient (ICC). This study focused on the reliability of the relative relationships among the measured values. To this end, a two-way mixed effects, single rater ICC (ICC (3,1)) was calculated using R studio version 4.2.2 (R Core Team, 2022) and the “psych” package (ver. 2.3.9). The reliability evaluation was based on 95% confidence intervals [35], and the outcomes revealed almost satisfactory matches (Table 3; S1 Fig.). Consequently, we used the measurements from the first author for subsequent analyses.

### Behavioral observations

We conducted two types of behavioral observations to investigate caretaking behaviors directed toward the target infants and examined the development of their motor skills. For recording caretaking behaviors, two female (Chonpe ‘01’21, Kusha ‘98’21) and two male infants (Cooper ‘94’02’21, and Cooper ‘94’01’21) were selected from the eight included in this study. These four individuals were born after the park-closure period and had minimal missing facial photograph data. We used focal animal sampling [36] between August and December 2021, with each session lasting 10–30 min, when the target infants were 12–23 weeks of age. This age corresponds to the period when infants are often separated from the mother and starts to actively engage with non-mothers. The indices focused on were the duration of affiliative physical contact with the mother or non-mother, including cuddling, grooming, carrying, and learning. The average observation duration per infant was 495.0 minutes (*SD* = 16.6).

Furthermore, we studied the developmental stages of all eight infants to record their motor skills and exploratory behaviors. The target behaviors were the five behavioral milestones linked to mobility and exploration in primates (Table 5). We recorded the presence or absence of these milestones in each subject while collecting facial photographs. Behavioral monitoring was initiated immediately after birth and persisted until each milestone was observed at least once in every infant.

**Table 5.** Description of the five behavioral milestones observed in Japanese macaque infant development.

| Behavioral milestones | Definition |
| --- | --- |
| Walk | Infants walk independently. |
| > 2 m away from the mother | Infants move > 2 m away from their mothers. |
| > 5 m away from the mother | Infants move > 5 m away from their mothers. |
| > 10 m away from the mother | Infants move > 10 m away from their mothers. |
| Social play | Infants play with other monkeys, including chasing and wrestling. |

### Data analysis

R Studio (version 4.2.2) was used for all statistical analyses. To prepare the data for analysis, individual Z-scores were computed for each facial feature. This involved subtracting the overall means from the individual measurements and dividing them by the standard deviations. The resulting Z-scores were used as the basis for subsequent statistical assessments.

### (1)#Identifying infantile facial features and quantifying infantile faces

First, to examine infantile facial features in Japanese macaques, linear mixed models using the “glmmTMB” package (ver. 1.1.3) were used to compare the Z-scores for each facial index across three age categories: infants, juveniles, and adults. The response variable was each individual’s facial index, the explanatory variables included age category and sex, and the random effect was individual ID. In order to verify the multicollinearity, we checked whether variance inflation factors (VIF) exceeded 3 [37] using the “performance” package (ver. 0.10.5) [38]. Next, using the Akaike information criterion (AIC), we compared the model fit between the model with all explanatory variables (full model) and that without them (null model). We selected the model with the smallest AIC value as the final model [39]. In cases where age exhibited a significant association with the explanatory variable, we conducted multiple comparisons by the Tukey method using the “multcomp” package (ver. 1.4.25).

Next, we defined infantile face scores (IFS) for Japanese macaques. This variable was referenced from Glocker et al. [8] and indicates the mean value of the Z-scores of all infantile facial features ascertained in this study. Prior to IFS calculation, sign reversal was applied to facial features with smaller Z-scores for infants. This methodology provided us with a valid means of quantifying infantile faces of Japanese macaques.

### (2)#Association between infantile faces and caretaking behaviors

To examine the link between infantile faces and caretaking behaviors toward infants, we performed generalized linear mixed models using the “glmmTMB” package. The explanatory variable was the total duration of affiliative contact with the mother or non-mother (in integer seconds) recorded on each observation day. The explanatory variables were the IFS, age (in days) on the day of observation, and sex. The IFS were the measured values of the photos taken on the day closest to the date of behavioral recording. Moreover, we introduced each day’s observation duration (in min) as an offset term. Because the response variables included numerous zeros (16/80 for mothers and 41/80 for non-mothers), we fitted four potential distributions as error structures: Poisson distribution, negative binomial distribution, zero-inflated Poisson distribution, and zero-inflated negative binomial distribution. Null models were also constructed for Poisson and negative binomial distributions. The model with the smallest AIC value was selected as the final model. To check multicollinearity, VIF was computed with the “performance” package (ver. 0.10.5). In cases where the highest 95% confidence interval of the VIF exceeded 3, we excluded the variable with the highest VIF, other than IFS, from the models.

### (3)#Association between infantile facial development and mobility development

Using the IFS, we explored the development of infantile faces within the first 24 weeks of age. To effectively examine the potential non-linear development of infantile faces, we formed three types of regression models and compared their fitting. The first was a linear model, the second was a linear model with the squared term for age as an additional explanatory variable, and the third was a generalized additive model (GAM). The smoothing parameter for the GAM was the generalized cross-validation method. In these models, IFS was the response variable and age (in days) was the explanatory variable. We used the “lm” function for the linear models and the “gam” function from the “mgcv” package (ver. 1.9.0) for GAM. Among the three candidate models and the null model, the model with the smallest AIC value was chosen as the final model for each individual. When multiple models demonstrated equal AIC values, the simpler model was selected as the final model.

Finally, to explore the associations between IFS development and infant mobility, we created figures showing the dates of the initial observation of five behavioral milestones and the date of the estimated IFS peak within the first 24 weeks of life [40]. Note that the behavioral milestone data only indicate the date of the first observation, and it is possible that the infants may have exhibited the behavior prior to our first records. Additionally, because of the study interruption between April and May 2021, continuous recordings were stopped for four infants (Ai’02’21, Cooper’90’01’21, Cooper’89’02’21, and Cooper’14’21; missing observations at 11–37, 0–18, 0–17, and 0–14 days of age, respectively). Therefore, we excluded parts of the data from these four individuals from the analysis. Specifically, we evaluated whether the age at which the corresponding behavioral milestone was initially observed in other infants overlapped with or was less than the unobservable age in each of the four infants. If such overlap or earlier records in other infants were found, the data on that milestone for that infant were excluded from the analysis. This led to the exclusion of four behavioral milestones (observed at least 2 m away, 5 m away, and 10 m away from the mother, and social play) for Ai’02’21 and two behavioral milestones (walking and being at least 2 m away from the mother) for Cooper’90’01’21, Cooper’89’02’21, and Cooper’14’21 from the analysis.

### Ethics statements

The study adhered to the Guidelines for Field Research established by the Ethics Committee of the Primate Research Institute, Kyoto University. The Arashiyama Monkey Park Iwatayama preapproved this study.

## Results

### Identifying infantile facial features and quantifying infantile faces

For all facial features, the AIC of the full model examining infantile features was smaller than that of the null model (S1 Table), resulting in the selection of the full model for all features. The VIF of the explanatory variables remained below 3 in all six full models, showing that there was no multicollinearity among the explanatory variables. For all facial features, age categories showed significant associations with the response variables (Table 6). Multiple comparisons revealed that in FWHL, FoLFaL, and EWFW, infants exhibited significantly higher Z-scores than both juveniles and adults (Fig 2). In contrast, the Z-scores of NLHL and NWFW were significantly lower in infants than in juveniles and adults (Fig 2). In the context of the MWFW, infants exhibited significantly lower Z-scores than adults and tended to have smaller Z-scores than juveniles (Fig 2). These findings indicate that infants showed significantly different values compared to adults and approximately distinct values compared to juveniles for all six indices. Consequently, we judged all six indicators as infantile facial features in Japanese macaques.

**Fig 2.**
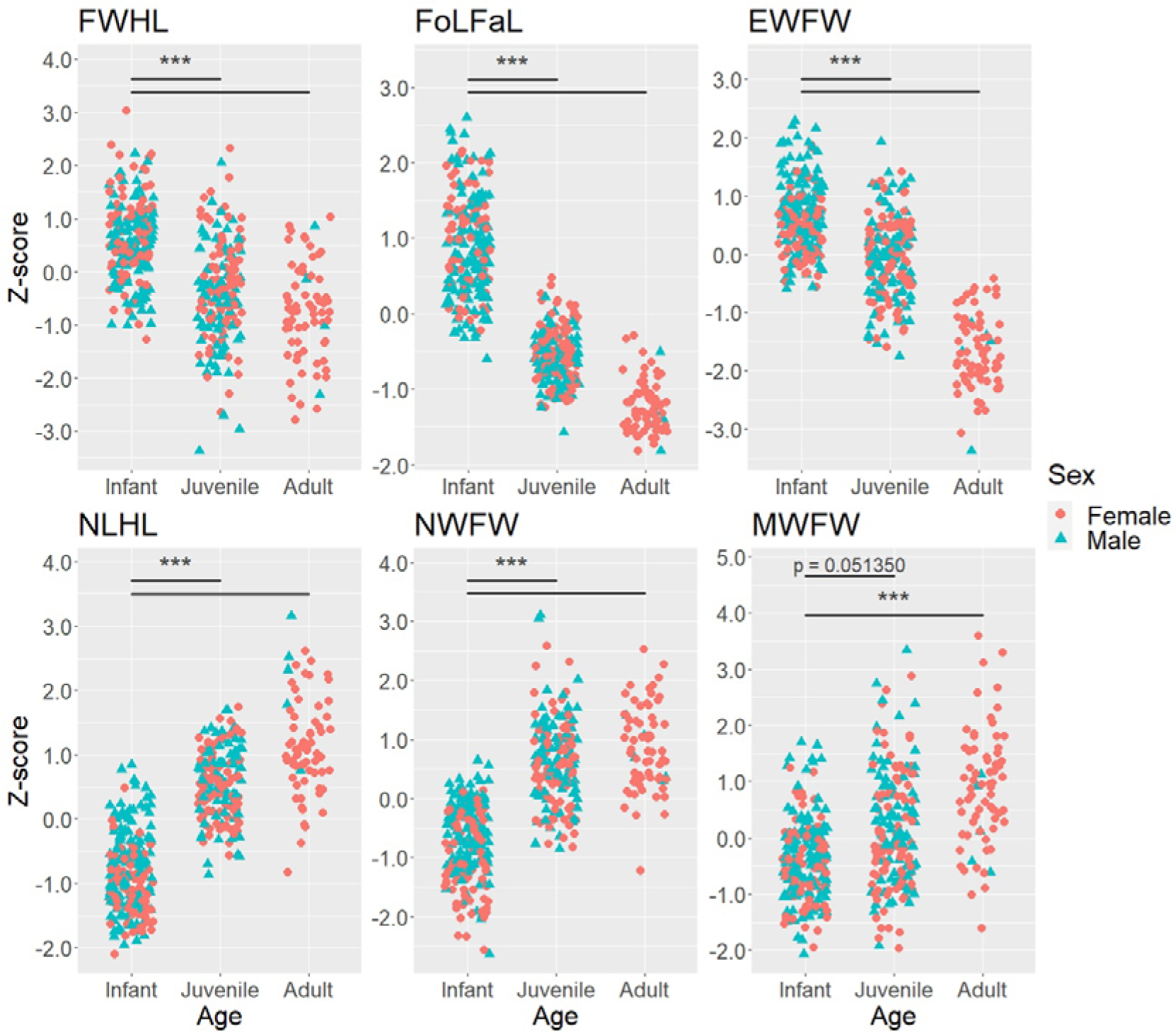
Comparisons of each facial feature between infant, juvenile, and adult Japanese macaques. “***” indicates p < 0.01.

**Table 6.**
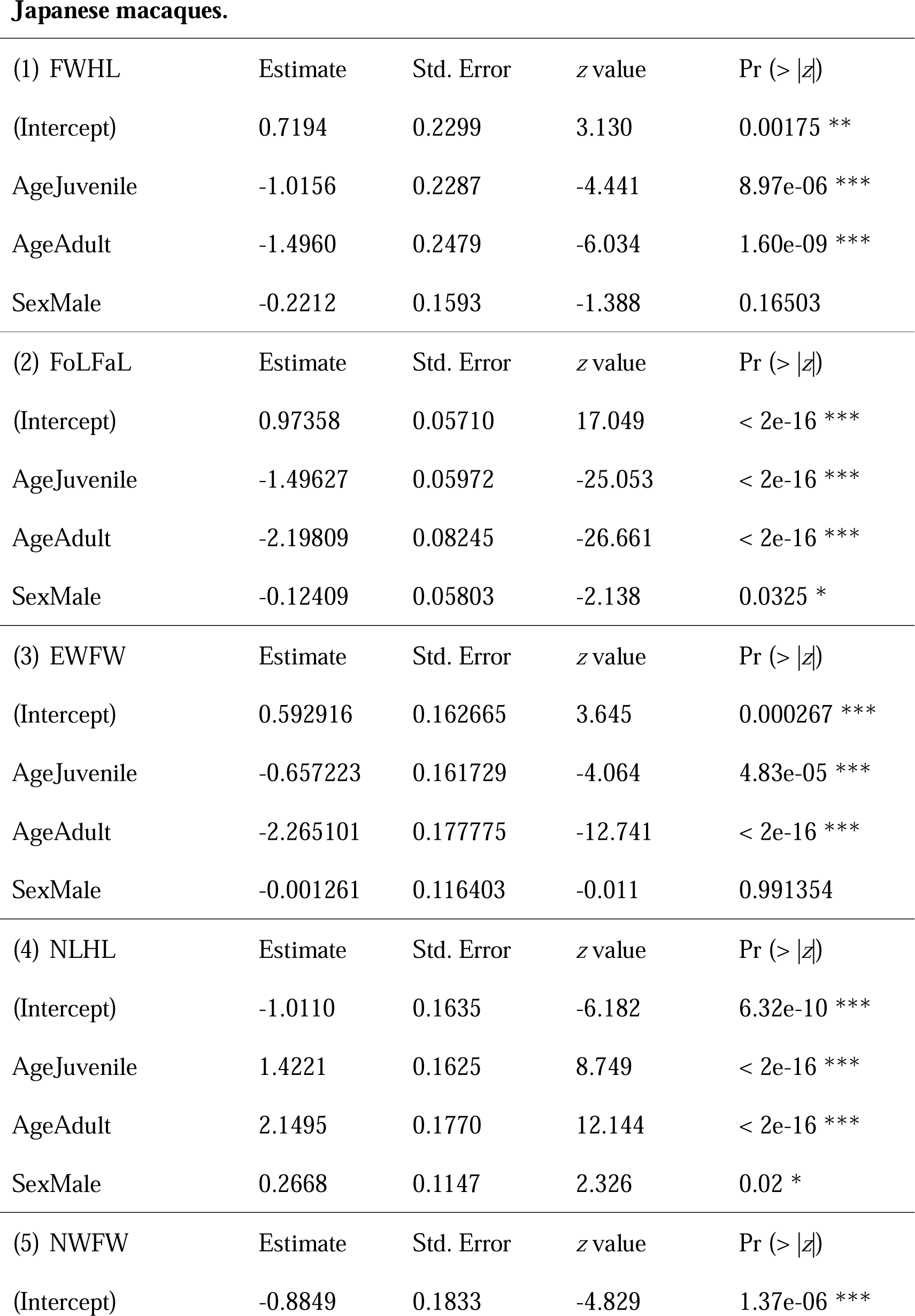
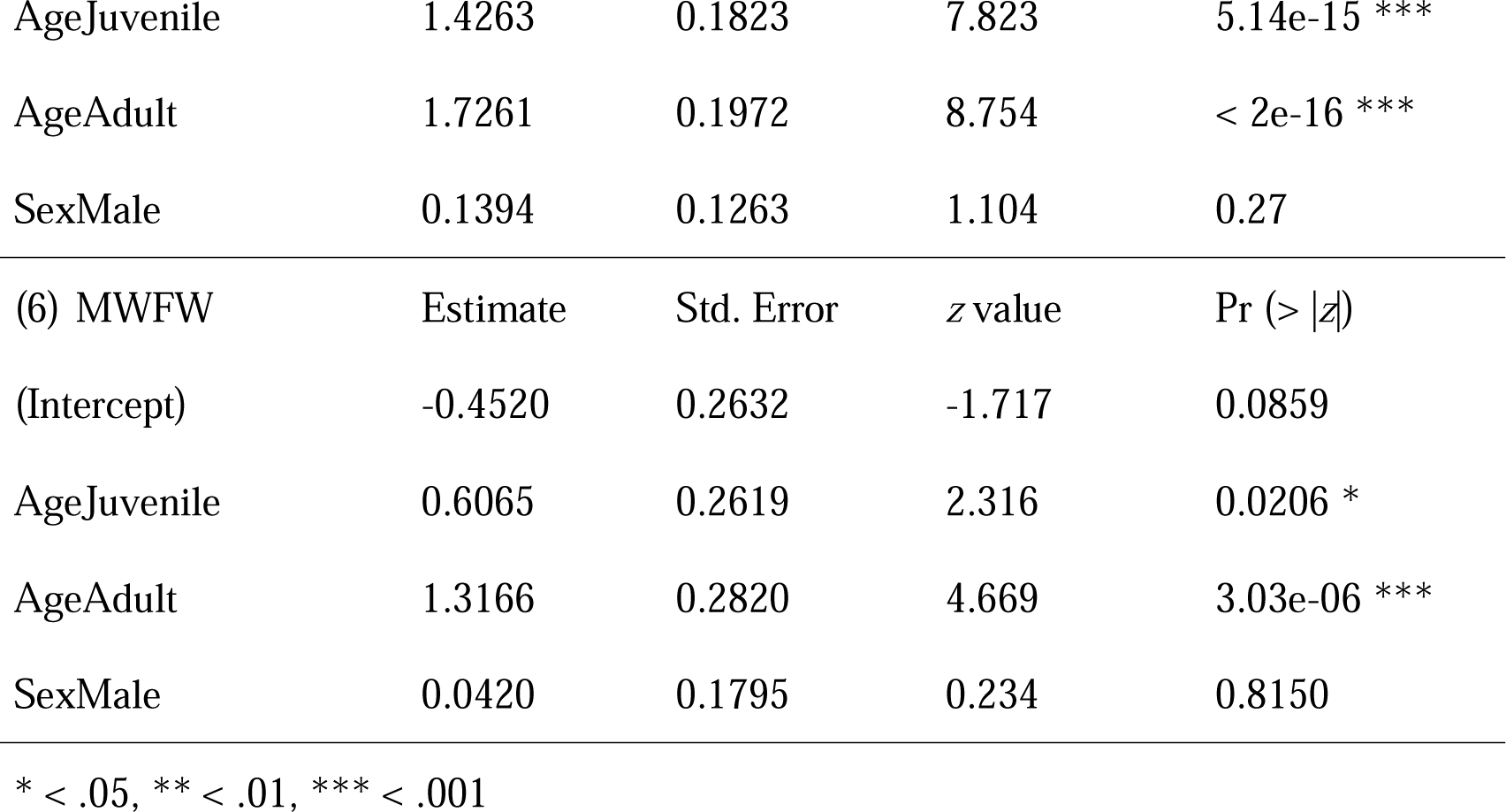
GLMM results comparing the facial index between age categories in Japanese macaques.

Based on these findings, IFS was defined as the mean value of FWHL, FoLFaL, EWFW, NLHL, NWFW, and MWFW. When calculating the IFS, we reversed the signs of NLHL, NWFW, and MWFW. When plotting IFS with age, it apparently decreased from early life to adulthood (Fig 3), indicating that IFS is a robust quantitative measure for capturing the infantile faces of Japanese macaques.

**Fig 3.**
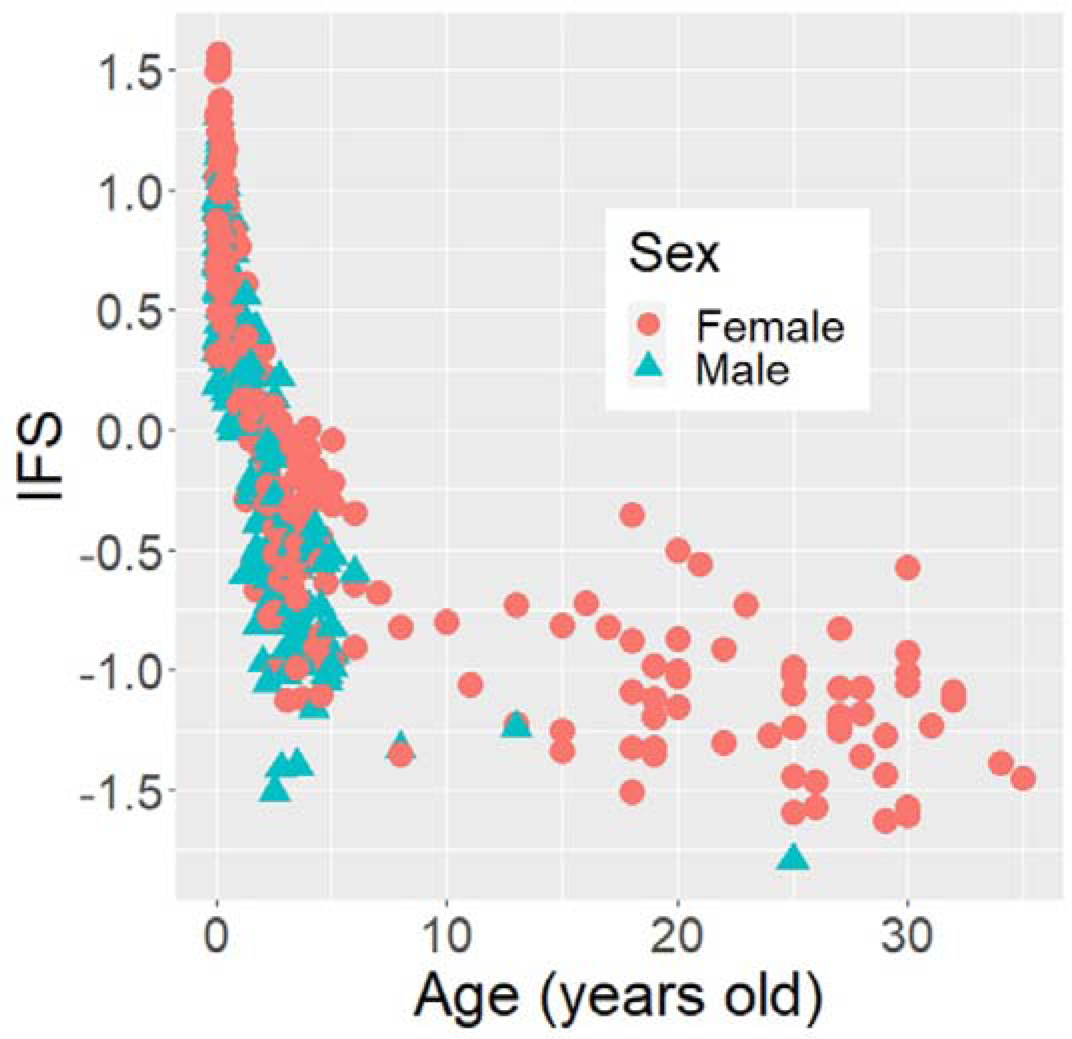
Developmental changes of IFS of Japanese macaques at all ages. The ages of adults were included as whole numbers (integers), while the ages of infants and juveniles were included as the age in days divided by 365 to allow direct comparisons with adults.

### Associations between infantile faces and caretaking behaviors

Among the models of the total duration of affiliative contact with the mother, the zero-inflated negative binomial model showed the smallest AIC value (S1 Table). However, the highest 95% confidence interval of the VIF for age was 5.03, indicating multicollinearity. Therefore, we formulated models with each error distribution, excluding age. Among them, the AIC was also the smallest in the model with the zero-inflated negative binomial distribution (S1 Table), and the VIF for this model was below 3. The results indicated the absence of a significant association between the IFS and the duration of affiliative contact with the mother (Table 7).

**Table 7.** GLMM results examining the association between IFS and affiliative contact duration with caregivers of Japanese macaque infants.

| (1) Affiliative contact duration with the mother |  |  |  |  |
| --- | --- | --- | --- | --- |
|  | Estimate | Std. Error | z value | Pr (> \|z\|) |
| (intercept) | 3.28446 | 0.40567 | 8.096 | 5.66e-16 *** |
| IFS | 0.39495 | 0.48319 | 0.817 | 0.414 |
| SexMale | 0.05267 | 0.24366 | 0.216 | 0.829 |
| (2) Affiliative contact duration with nonmothers |  |  |  |  |
|  | Estimate | Std. Error | z value | Pr (> \|z\|) |
| (intercept) | 1.99018 | 0.42430 | 4.69 | 2.73e-06 *** |
| IFS | 0.01227 | 0.60547 | 0.02 | 0.984 |

In the model examining the total duration of affiliative contact with non-mothers, the zero-inflated negative binomial model showed the smallest AIC value (S1 Table). However, the highest 95% confidence interval of VIF for IFS and sex were 5.94 and 5.74, respectively. Thus, sex variables were excluded from each model. Subsequently, the negative binomial model was again selected by AIC (S1 Table), but the highest 95% confidence interval of VIF for IFS and age was very high (1.83 × 10^5^). Therefore, age was excluded from our model. Finally, among the models with only the IFS as an explanatory variable, the model with a zero-inflated negative binomial distribution displayed the smallest AIC value (S1 Table). The results demonstrated that the IFS was not significantly associated with the duration of affiliative contact with non-mothers (Table 7).

### Development of infantile faces and behavioral associations

After formulating four candidate models, the AIC selected the linear model for Cooper’90’01’21 and Kusha’98’21, GAM for Cooper’94’02’21, Ai’02’21, Cooper’14’21, Cooper’94’01’21, and Cooper’89’02’21, and the null model for Chonpe’01’21 (S1 Table). Based on these outcomes, we found two distinct patterns of IFS development (Fig 4). Three infants (Cooper’90’01’21, Kusha’98’21, and Cooper’94’02’21) showed consistent decreases in IFS until 24 weeks of age. Conversely, the IFS of four individuals (Ai’02’21, Cooper’14’21, Cooper’94’01’21, and Cooper’89’02’21) increased immediately after birth and then decreased, although the 95% confidence intervals were wide. The peak IFS estimates for these four infants were 41.22, 45.71, 66.17, and 72.18 days, respectively. Notably, the three infants with consistently decreasing IFS included two females and one male, whereas all four infants with a hump-shaped IFS development were male (Fig 4).

**Fig 4.**
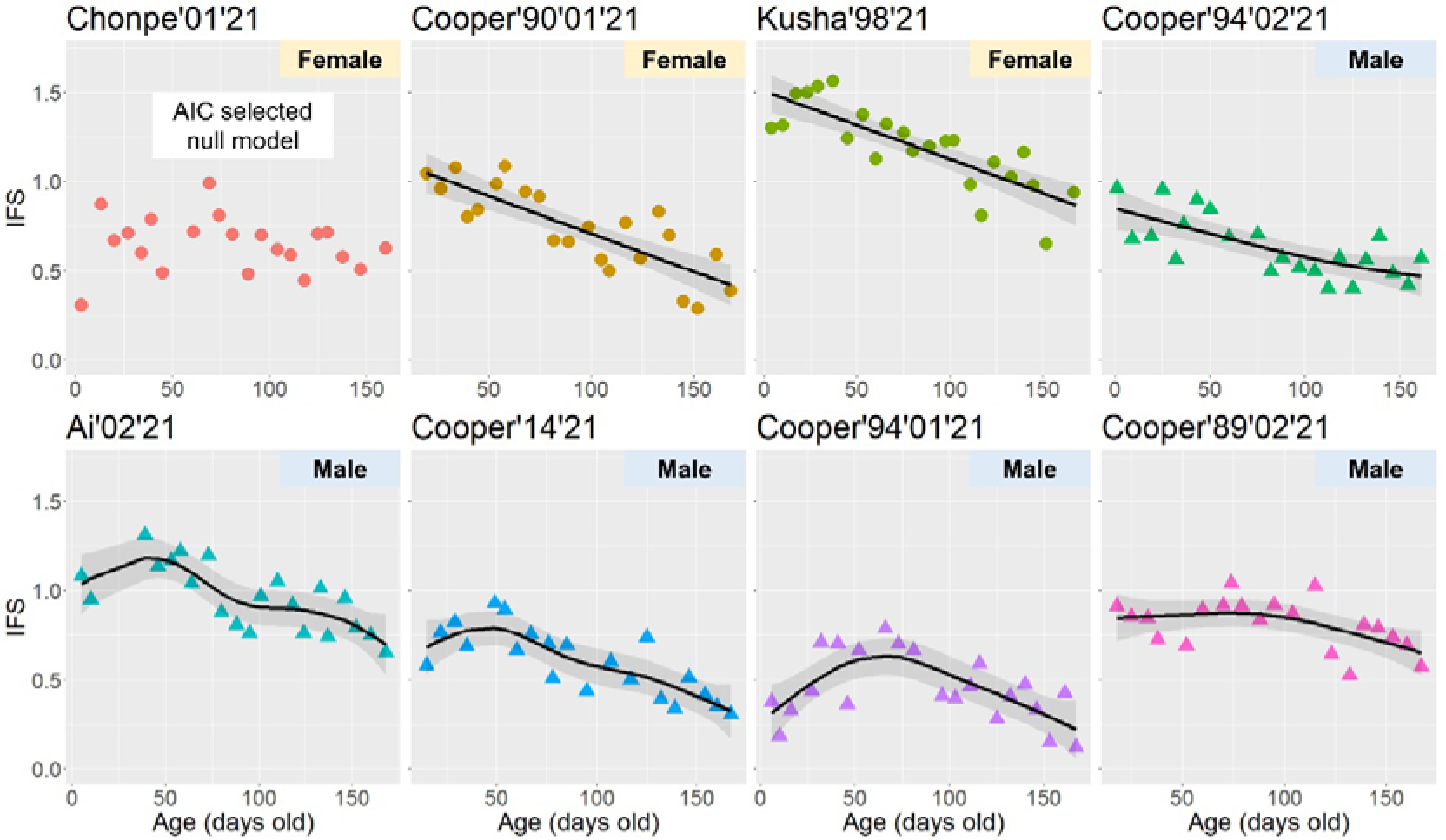
Individual differences in IFS development of Japanese macaque infants within the first 24 weeks of life. Black lines and gray ribbons represent the estimated values and 95% confidential intervals. The circle and triangle plots indicate females and males, respectively.

Fig 5 depicts the association between the age of the estimated IFS peak for each infant and the behavioral milestones. We found no consistent trend between the IFS and infant mobility or exploratory behavior.

**Fig 5.**
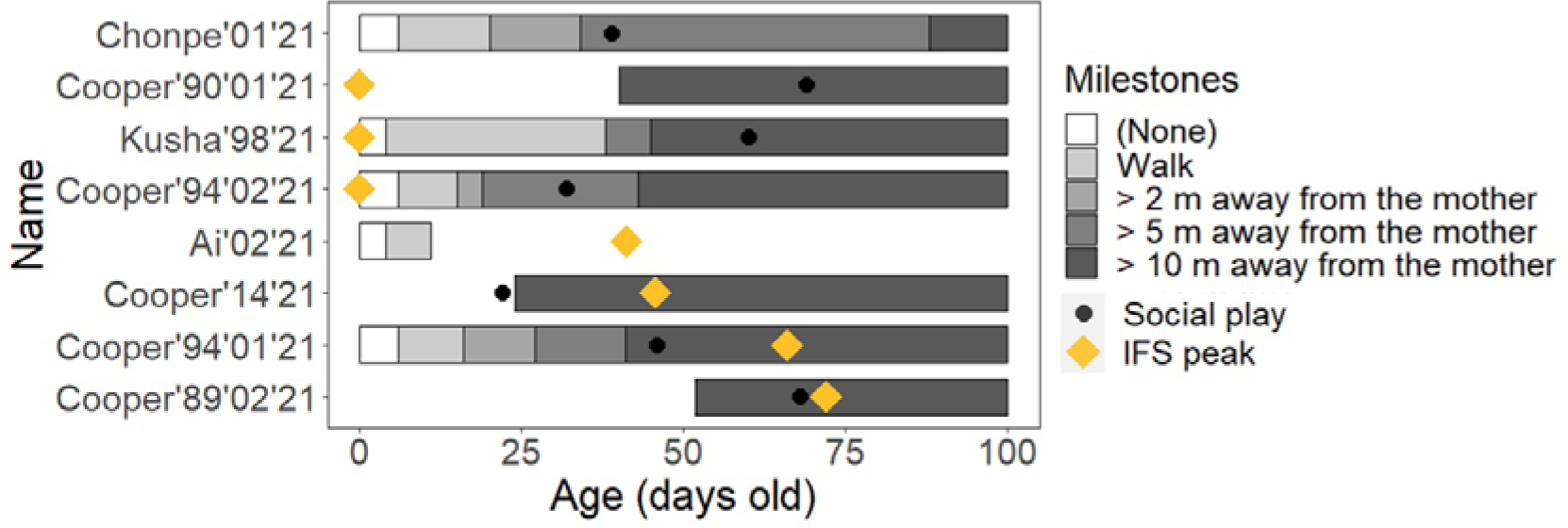
The age with estimated IFS peaks and the first-observed age in days of the five behavioral milestones in the eight Japanese macaque infants observed. Certain information remains absent due to the temporary suspension of observations.

## Discussion

### Infantile facial features and their association with caretaking behaviors

The first aim of this study was to ascertain the infantile facial features of Japanese macaques. The findings revealed significant differences between the infants and adults across all six indices. Moreover, a consistent trend was observed in Japanese macaques and humans, characterized by larger face width, forehead length, and eye width, along with a smaller nose and mouth. Therefore, Japanese macaques have these six indices as infantile facial features, similar to humans. Some features, such as face width, forehead length, and eye width, have also been reported as shared infantile features among great apes [23], suggesting the likelihood that these infantile facial features are shared at least among Catarrhini.

We also found differences between the infantile facial features of Japanese macaques compared to great apes. Kawaguchi et al. [23] deduced that the sizes of the nose and mouth were not shared infantile features among great apes. In contrast, we found that nose and mouth sizes were associated with infantile features in Japanese macaques. The differences between these studies could be attributed to two factors. First, there may be species differences in infantile facial features [23, 24, 41]. For example, infant chimpanzees have rounded supraorbital torus as infantile features that have not yet been reported in humans [24]. Differences in the nose and mouth size between great apes and Japanese macaques may also reflect interspecific differences. If so, the interspecies variation in infantile features may surpass what Lorenz [7] initially assumed. Second, the methodological differences between our study and Kawaguchi et al. [23] may have contributed to this variation. This study directly measured candidate facial features, whereas Kawaguchi et al. [23] used geometric morphometrics, which treated facial landmarks as coordinates and extracted principal components to describe the overall facial morphology. This methodological difference potentially accounts for the heightened sensitivity to the nose and mouth size in our study. Future studies should consider these methodological differences to extend the research to a broader range of species to reveal the interspecific similarities and variations in infantile facial features.

The second objective was to examine whether infantile faces in Japanese macaques exhibit a notable link with caretaking behaviors toward infants. We did not find significant associations between infantile faces and affiliative contact with mothers or non-mothers. This finding is consistent with that of Koda et al. [28], who reported no preference for infant faces in this species. Additionally, when exploring the responses to infantile faces in some great apes, chimpanzees displayed a significant but very weak preference, and bonobos exhibited none [24, 27]. In contrast, infantile faces in humans can elicit strong responses and may facilitate caretaking motivation and affectionate care [8–11, 17, 18]. Contrary to Lorenz [7], these results suggest significant species differences in the link between infantile facial shape and caretaking behaviors toward infants, with a much stronger link in humans.

The underlying factors contributing to the heightened reactivity in humans to infantile faces remain unclear, but several possibilities exist [42, 43]. For example, Glocker et al. [17] emphasized the importance of a higher frequency of non-maternal caretaking behaviors. In cooperative breeding species like humans [44], nonmother’s preference for shared infantile facial features within the species may provide adaptive advantages to them. To test this hypothesis, it is imperative to examine how other cooperative breeding primates, such as Callithrichidae [45–47], respond to infantile facial features. As another possibility, interactions with other infantile physical features, such as the body or fur color specific to infants (known as “infantile coloration”), may also hold significance. Chimpanzees exhibit a heightened response to infantile facial skin color compared to facial shape [24], whereas humans lacking distinct infant coloration [43] display a stronger response to facial morphology [8–11]. This suggests a trade-off between increased reactivity to infantile coloration and stronger reactions to infantile facial morphology. However, this interpretation is inconsistent with the finding that bonobos, which lack infantile coloration, did not prefer infant images to adult ones [27]. Further comparative studies are warranted to determine whether and how infantile coloration affects reactivity to infantile facial shape among primates.

Note that the response of Japanese macaques to infantile faces remains a topic without conclusive evidence. To the best of our knowledge, Koda et al. [28] and the present study are the only empirical investigations exploring the connection between infant faces and the responses of Japanese macaques. However, both studies were conducted with limited sample sizes, consisting of only two nulliparous females in Koda et al. [28] and four infants in the current study. Additionally, the behavioral observations employed in our study were simplified; we could not target potentially associated behaviors other than affiliative contact and direct care, such as the interest in infants [48–50] or careful behavioral responses [18, 51, 52]. Furthermore, we could not control for other potential factors affecting caretaking behavior in Japanese macaques, such as maternal age, parity [53, 54], and social relationships between non-maternal caregivers and mothers [55]. Therefore, future research should include more precise observational and statistical designs to further examine the responses to infantile faces among primates.

### Development of infantile faces and associations with mobility

The third objective was to investigate the development of infantile faces in Japanese macaques during the early postnatal period. Although studies on non-human primates have explored the developmental process of infantile coloration during infancy [56, 57], this study represents the first empirical example of the development of infantile faces and their individual differences in infant primates. Our findings revealed that infantile faces did not necessarily reach their peak proportions immediately after birth but instead peaked between 40 and 70 days of age in some subjects. This hump-shaped developmental trajectory of infantile faces parallels findings in humans [15, 20, 21, 22]. However, since we could only access the averaged results from previous studies in humans, it remains uncertain whether human infants also have individual differences in infantile facial development or whether the hump-shaped developmental trajectory is more commonly observed in humans. Research on primates, including humans, is essential for understanding the variations in the development of infantile faces within and across species.

The current study was unable to identify the factors responsible for individual differences in facial development in macaque infants. However, there are several potential explanations. The first was the infant sex. In this study, four of the five male infants exhibited a hump-shaped facial development pattern, whereas no female infants did. This suggests that the hump-shaped development of infantile faces is a characteristic of males, in macaques at least. While sex differences in the cranium of young Japanese macaques have been explored, the examination of neonates was constrained by the unavailability of female infant specimens [58]. Therefore, it is necessary to investigate sex differences in the morphological development of infant Japanese macaque faces in greater detail. Another possible factor is the health status of the infants. In humans, infant health conditions may manifest as changes in facial appearance [59]. If a similar phenomenon was observed in Japanese macaques, the development of infantile faces could be partly affected by early postnatal health. Given that our study did not record detailed behavioral data of infants in the early postnatal period, this aspect warrants further exploration in future studies.

The fourth purpose of our study was to investigate whether infantile faces in Japanese macaques reach their peak at the onset of exploratory behavior. The hump-shaped development of infant faces might be linked to the function of capturing and retaining caregivers’ attention [15]. If infantile faces in non-human animal species also promote attention or caretaking behaviors, it would be valuable to examine this hypothesis in these species too. The current study did not demonstrate clear associations between peak IFS and the development of exploratory behaviors in Japanese macaques, although we found humped-shaped development of infantile faces in male individuals. This result was reasonable, given that infantile faces were not significantly associated with caretaking behaviors. Even in humans, the association between facial development and infant behaviors remains almost untested. Therefore, further research is warranted to examine the relationship between the development of infantile faces and infant behavior.

### Future directions

This study demonstrates that infantile facial features and the developmental patterns of infantile faces in Japanese macaques may exhibit trends similar to those of humans or non-human great apes, while also revealing partial species differences. Furthermore, we did not find any significant associations between infantile faces and behavioral indicators, as is the case in humans. Consistent with previous studies, these results suggest that although many morphological aspects of facial infantile-ness may be common across primates [7, 23], there are interesting interspecific variations in responses to these features, with particularly strong responsiveness in humans [8, 25, 27, 28]. Future research should target a broader range of lineages to further elucidate both the commonalities and variations in infant faces and the responses toward them.

Finally, as a prospective avenue for primate research, we emphasized the significance of physical features beyond frontal faces, such as profile, whole body, and body color. Although previous studies focused on frontal faces, Lorenz [7] originally suggested the role of other physical features in eliciting care among animals. Indeed, early studies in humans have indicated connections between infantile profiles or full-body images and perceptions of cuteness by observers [9, 10, 60]. Japanese macaques also demonstrate a preference for full-body images of conspecific infants [61]. In addition, many primates exhibit specific infantile coloration [43, 62], which may stimulate adult interest and caretaking behaviors [16, 24, 62–65]. In real-world caretaking contexts, caregivers have abundant exposure to various infantile physical features that are not limited to faces alone. Therefore, future research should include physical features beyond frontal facial characteristics to enable more detailed comparative studies of infantile physical features and the responses to them.

## Acknowledgments

We thank Shinsuke Asaba and the staff at Arashiyama Monkey Park Iwatayama for their invaluable support in facilitating our research. We are grateful to Yuri Kawaguchi for her invaluable advice in designing this study. We acknowledge the contributions of Natsu Mizuno, who helped with the reliability assessment of our measurements. This study was supported by the Collaborative Research Program of Wildlife Research Center, Kyoto University, and the Global Education Office in Graduate School of Education, Kyoto University.

## Author contributions

Conceptualization: Toshiki Minami.

Formal analysis: Toshiki Minami, Takeshi Furuichi. Investigation: Toshiki Minami.

Methodology: Toshiki Minami. Supervision: Takeshi Furuichi.

Visualization: Toshiki Minami.

Writing – original draft: Toshiki Minami. Writing – review & editing: Takeshi Furuichi.

## Data availability statement

All data utilized in the analysis are within the Supporting Information.

## Supporting information

S1 Fig. Plots of the measurements of nine Japanese macaque facial parts from 118 photographs measured by two raters to assess inter-rater reliability.

S1 Table. AIC results for the model selection in this study. S1 File. Raw data files for all analyses in this study.

