## Supplementary material for "Examining infantile facial features and their influence on caretaking behaviors in free-ranging Japanese macaques (*Macaca fuscata*)": S1 Fig & S1 Table

Research Article

Short title: Infantile facial features and their behavioral effects in Japanese macaques.

Toshiki Minami^1,2*^, Takeshi Furuichi^1,3^

^1^ Primate Research Institute, Kyoto University, Inuyama, Aichi, Japan

^2^ Graduate School of Education, Kyoto University, Kyoto, Kyoto, Japan

^3^ Wildlife Research Center, Kyoto University, Kyoto, Kyoto, Japan

^*^ Corresponding author

ORCID: 0000-0001-5476-3896

S1 Fig. Plots of the measurements of nine Japanese macaque facial parts from 118 photographs measured by two raters to assess inter-rater reliability.

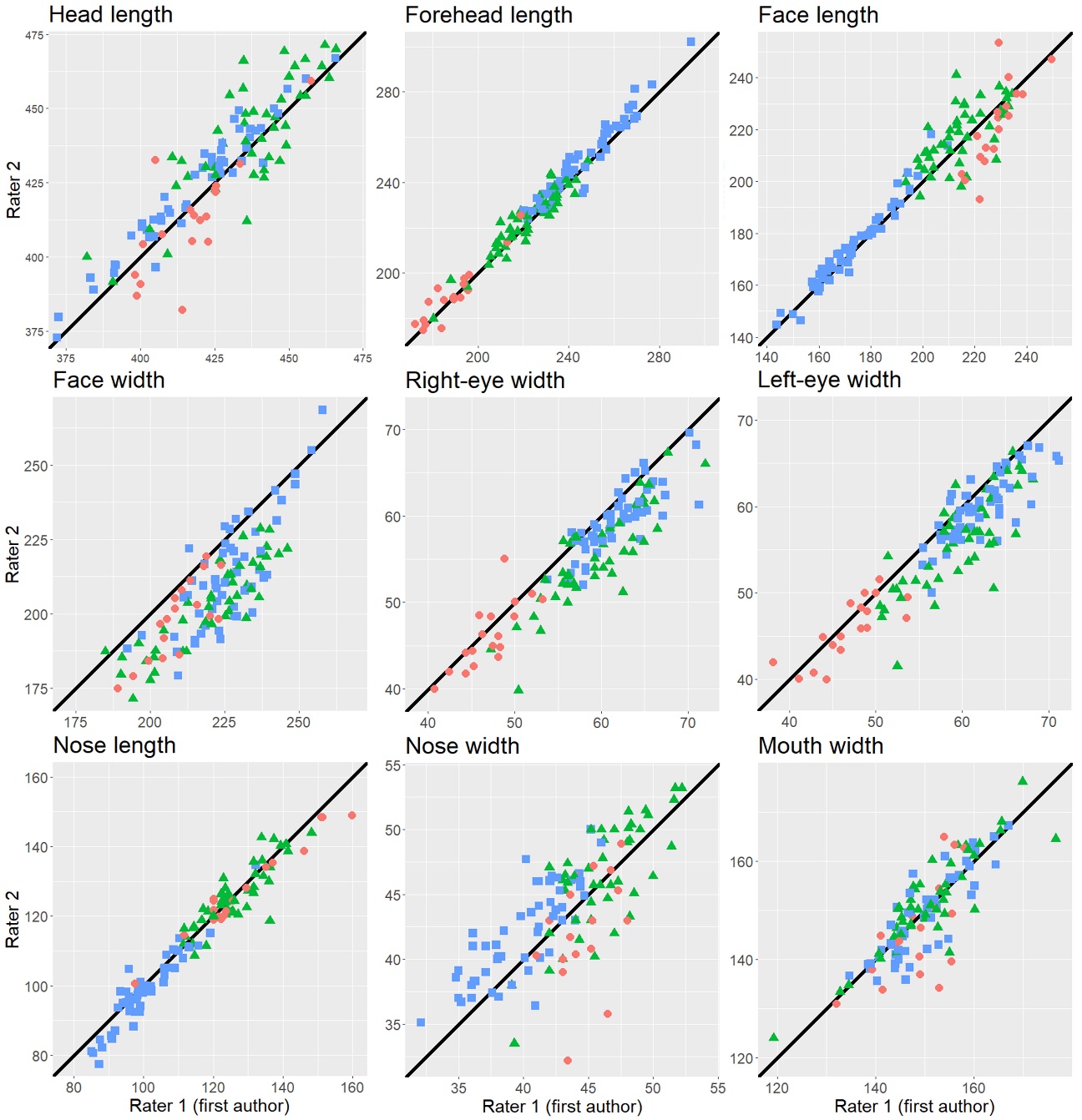

Note) The color-coded shapes, such as orange circles, green triangles, and blue squares, correspond to adult individuals, juveniles, and infants, respectively. The black lines depict instances where the ratings given by two raters precisely matched (y = x).

S1 Table. AIC results for the model selection in this study.

| (1) Identifying infantile facial features | | | | | | | | | |
| --- | --- | --- | --- | --- | --- | --- | --- | --- | --- |
|  | Null model | | | | Full model | | | | |
| EWFW | 1100.117 | | | | **1075.59** | | | | |
| FoLFaL | 906.9761 | | | | **724.5124** | | | | |
| EWFW | 1008.304 | | | | **858.3772** | | | | |
| NLHL | 869.6228 | | | | **788.3585** | | | | |
| NWFW | 888.7603 | | | | **839.387** | | | | |
| MWFW | 1160.364 | | | | **1137.672** | | | | |
| (2) Associations between infantile faces and caretaking behaviors^*^ | | | | | | | | | |
|  | Null model for Poisson | Poisson | | Zero-inflated Poisson | | Null model for negative binomial | Negative binomial | | Zero-inflated negative binomial |
| Affiliative contact duration with the mother (including all explanatory variables) | 33830.46 | 31396.95 | | 14656.86 | | 978.8119 | 983.8066 | | **915.8777** |
| Affiliative contact duration with the mother (after excluding the age variable) | 33830.46 | 31724.97 | | 14902.92 | | 978.8119 | 981.8849 | | **913.9064** |
| Affiliative contact duration with nonmothers (including all explanatory variables) | 18399.7 | 18080.63 | | 8475.734 | | 550.4012 | 555.9225 | | **541.5085** |
| Affiliative contact duration with nonmothers (before excluding the age variable) | 18399.7 | 18400.47 | | 8585.18 | | 550.4012 | 553.9227 | | **541.7889** |
| Affiliative contact duration with nonmothers (after excluding the age variable) | 18399.7 | 18111.02 | | 8672.398 | | 550.4012 | 552.3985 | | **540.4813** |
| (3) Development of infantile faces | | | | | | | | | |
|  | Null model | | Linear model | | Linear model with squared term | | | GAM | |
| Chonpe’01’21 | **-15.90611** | | -14.22361 | | -14.37325 | | | -14.97141 | |
| Cooper’90’01’21 | 2.11505 | | **-20.28548** | | -18.32175 | | | -20.28548 | |
| Kusha’98’21 | 0.5979557 | | **-22.40326** | | -21.14992 | | | -22.40326 | |
| Cooper’94’02’21 | -12.62248 | | -26.14362 | | -25.54728 | | | **-26.41706** | |
| Ai’02’21 | -8.419618 | | -19.53832 | | -19.3766 | | | **-24.09722** | |
| Cooper’14’21 | -7.988981 | | -23.87437 | | -24.45388 | | | **-26.28746** | |
| Cooper’94’01’21 | -7.558351 | | -7.731556 | | -18.53666 | | | **-20.47128** | |
| Cooper’89’02’21 | -19.39724 | | -21.90838 | | -24.08174 | | | **-24.57192** | |
| The numbers in bold are the lowest AIC among the comparing models. When some AICs were equal, simpler models were selected.  ^*^ AICs in the models after excluding explanatory variables based on VIF. | | | | | | | | | |
